# Male-biased stone tool use by wild white-faced capuchins (*Cebus capucinus imitator*)

**DOI:** 10.1101/2023.09.04.556228

**Authors:** Zoë Goldsborough, Margaret C. Crofoot, Brendan J. Barrett

## Abstract

Tool-using primates often show sex differences in both the frequency and efficiency of tool use. In species with sex-biased dispersal, such within-group variation likely shapes patterns of cultural transmission of tool-use traditions between groups. On the Panamanian islands of Jicarón and Coiba, a population of white-faced capuchins (*Cebus capucinus imitator)*—some of which engage in habitual stone tool use—provide an opportunity to test hypotheses about why such sex-biases arise. On Jicarón, we have only observed males engaging in stone tool use, whereas on Coiba, both sexes are known to use tools. Using 5 years of camera trap data, we show that this variation reflects a true sex difference in tool use rather than a sampling artefact, and then test hypotheses about the factors driving this pattern. Differences in physical ability or risk-aversion, and competition over access to anvils do not account for the sex-differences in tool-use we observe. Our data show that females are physically capable of stone tool use: females on Coiba and juveniles on Jicarón as small and smaller than adult females regularly engage in tool use. Females also have ample opportunity to use tools: the sexes are equally terrestrial, and competition over anvils is low. Finally, females rarely scrounge on left-over food items either during or after tool-using events, suggesting they are not being provisioned by males. Although it remains unclear why white-faced capuchin females on Jicarón do not use stone-tools, our results illustrate that such sex biases in socially learned behaviors can arise even in the absence of obvious physical, environmental and social constraints. This suggests that a much more nuanced understanding of the differences in social structure, diet and dispersal patterns are needed to explain why sex-biases in tool use arise in some populations but not in others.

## Introduction

A small but taxonomically diverse set of primate species are known to use tools for extractive foraging (e.g., Burmese long-tailed macaques [*Macaca fascicularis*], Gumert et al., 2009); tufted capuchins [*Sapajus apella*], Ottoni & Mannu, 2001; chimpanzees [*Pan troglodytes*], Goodall, 1986; white-faced capuchins [*Cebus capucinus imitator*], Panger et al., 2002). Despite providing access to high-quality food items that are often otherwise unavailable, tool-use behavior tends to be highly variable, meaning that only some members of a tool-using group or population actually use tools. Sex-differences in extractive foraging behavior are often observed in tool-using primates (Falótico et al., 2021; Gumert et al., 2011; Lonsdorf, 2017). In the majority of instances, such biases are a matter of degree not kind: one sex might use tools more frequently or with more proficiency than the other, but both sexes show the tool use behavior. Identifying the factors that drive sex differences in tool use is key to understanding how and why tool use traditions first arose, as well as the mechanisms responsible for their maintenance and patterns of transmission. Because tool use in primates spreads between individuals via social learning (Coelho et al., 2015; Krutzen et al., 2005; Lamon et al., 2017; Wild et al., 2019), and many species show sex-biased dispersal (Clutton-Brock & Lukas, 2012), differences in tool use frequency and proficiency between sexes has important implications for the cultural transmission of tool use behavior between groups.

Considerable attention has been paid to sex differences in tool use in primates, in part because of the hypothesized connection between such behavioral differentiation and the emergence of a sexual division of labor that is so prominent in human societies (Kuhn & Stiner, 2006). However, sex-differences in tool-use are not universal. Among the apes, orangutans (*Pongo pygmaeus*) and gorillas (*Gorilla gorilla*), show no reported sex differences in tool use (Breuer et al., 2005; Fox et al., 2004), although gorillas rarely show tool use in the wild. Tool-using bird species, for example, New Caledonian crows (*Corvus moneduloides*; Kenward et al., 2004) and woodpecker finches (*Cactospiza pallida*; Tebbich et al., 2002) show little evidence of sex-biased tool use. Where sex differences in tool use by animals do exist, neither males or females consistently emerge as the more frequent or proficient tool users. For example, while sponging behavior in bottle-nosed dolphins (*Tursiops sp.*) is female biased (Mann et al., 2012), tool use proficiency in captive Goffin cockatoos *(Cacatua goffini*) appears to potentially be higher in males (Auersperg et al., 2014) This variability suggests that different social and/or ecological pressures underly the patterns of sex-biased tool use we observe across the animal kingdom, raising the question: when and why do sex-biases in tool use emerge?

Energetic requirements—and specifically the high costs of pregnancy and lactation— are hypothesized to play a key role in driving sex-differences in tool-use among primates, although they have been invoked to explain both male- and female-biased patterns of tool-use. To explain female-biased tool use in primates, researchers have invoked the high nutritional demands of pregnancy and lactation which, they argue, lead to females investing more heavily in learning to use tools that increase their foraging efficiency (Lonsdorf, 2017; McGrew, 1992). Direct evidence supporting this hypothesis, however, is lacking. Female-biased tool use is known to exist in chimpanzees (*Pan troglodytes)* and bonobos (*Pan paniscus),* where females acquire skills faster (Boose et al., 2013; Lonsdorf et al., 2004), and use tools more frequently and, in some cases, more efficiently than males (Boesch & Boesch, 1981; Boose et al., 2013; Gruber et al., 2010), but these differences have not been explicitly tied to the energetic needs of females. Incongruously, it has also been speculated that the increased energetic demands of pregnancy and lactation might also give rise to male-biased tool use, since males can better sustain the energetic costs of high risk/high reward foraging strategies than females (de A. Moura & Lee, 2010). Among primates, male-biased tool use is found in the *Sapajus* and *Cebus* capuchin monkeys. In bearded capuchins (*Sapajus libidinosus*) and tufted capuchins (*Sapajus apella*) males use stone tools (Ottoni & Mannu, 2001; Spagnoletti et al., 2011; Visalberghi, 1987) more frequently than females and only males use probing stick tools (de A. Moura & Lee, 2010; Falótico & Ottoni, 2014). In blonde capuchins (*Cebus flavius*), termite fishing appears male-biased (Souto et al., 2011). However, there is currently no direct evidence linking the male-bias in *Sapajus* tool use to the energetic costs of failure. Rather, the male bias in *Sapajus* stone tool use is thought to be due to physical limitations of females, as males open more high resistance nuts than females (Spagnoletti et al., 2011). Further, sex differences in the male-biased probe tool-use behavior of bearded capuchins arise only later in life: while juveniles of both sexes engage in manipulation of sticks, only males develop the probing tool use, which is hypothesized to be the result of a sex motivation bias (Falótico et al., 2021).

In addition to differences in nutritional needs, morphological differences can also drive variation in behavior between sexes. In species with marked sexual dimorphism, which includes many primate species (Plavcan, 2001), the larger sex tends to be more efficient at behaviors that require significant strength. Male-biased nut-cracking by bearded capuchins likely arises as a result of this dynamic. Bearded capuchins show strong sexual dimorphism; female capuchins weigh only 60% as much as males and handle the large stones used for nut-cracking less efficiently (Fragaszy et al., 2010; Spagnoletti et al., 2011). Burmese long-tailed macaques (*Macaca fascicularis aurea)*, which use stone tool use to open coastal resources, provide additional evidence that differences in morphology underly sex-differences in tool-use behavior. Although similar to bearded capuchins in their degree of sexual dimorphism, male and female Burmese long-tailed macaques did not differ in tool use proficiency, but females used smaller hammerstones and targeted more sessile prey than males (Gumert et al., 2011). Marked sexual dimorphism may thus give rise to sex-based differences in tool-use behavior because the smaller sex is physically incapable of using tools, is inefficient enough that the costs of tool-use outweigh the benefits, or adopt alternate behavioral strategies, preferentially selecting different tools or focusing on alternate food resources.

Sex differences in behavior do not necessarily arise from differences in morphology; significant evidence suggests that male and female primates also differ in fundamental behavioral tendencies (Lonsdorf, 2017). A meta-analysis found that male primates show higher innovation rates than females (Reader & Laland, 2001), although no difference between sexes was found in a long-term study of a large population of white-faced capuchin monkeys (*Cebus capucinus*; Perry et al., 2017). Higher innovativeness of one sex can be driven by inherent cognitive differences, but may also be a result of sexes facing different constraints and needs in their environment. One sex may be at greater risk of predation, be more risk-averse or engage in more anti-predatory behaviors. For example, female chimpanzees spent more time building nests than males (Stewart & Pruetz, 2020) and female and juvenile chimpanzees built their nests at higher, more peripheral locations, potentially to decrease risk of predation (Stewart & Pruetz, 2013). While in woolly monkeys (*Lagothrix lagotricha poeppigii*), males engage in more vigilance behavior than females (Di Fiore, 2002). Sex differences in predation risk and risk perception can also lead to variation in tool use behavior. In the case of bearded and tufted capuchins, for example, females are thought to be more vulnerable on the ground where nut-cracking takes place, which may lead them to use tools less frequently than males (Fragaszy, 1990). Additionally, increased terrestriality correlates with the diversity of tool use repertoires in bearded capuchins (Falótico & Ottoni, 2023), suggesting sex differences in time spent on the ground could be a driver in variation between sexes in tool use behavior. While a study that explicitly tested key assumptions of the risk hypothesis found that female and male bearded capuchins did not differ in the amount of time they spent on the ground (de A. Moura & Lee, 2010), predation risk may still differ between the sexes and between sites.

Dominance and competition also affect the opportunities of males and females to access resources, perhaps including the sites and materials for tool-use. In white-faced capuchins, high ranking group members often displace lower ranking individuals at important food sites and, as a result, energy intake rates vary depending on rank (Vogel, 2005). Competition may be particularly strong when tool use relies on fixed, monopolizable resources, such as anvils, where high-ranking individuals can displace lower ranking individuals for access (e.g., in bearded capuchins, Spagnoletti et al., 2011; Verderane et al., 2013). When one sex is dominant over the other sex, the lower-ranking sex may have limited opportunities to use tools and thus engage in this behavior less often.

Sex-differences in tool use behavior can also arise as a result of sexual selection if, for instance, females favor inventive males or if males display their quality and strength through tool use (de A. Moura & Lee, 2010). For example, female bearded capuchins, who use stone and stick tools during foraging, were found to throw these stones and sticks at males as a part of sexual displays (Falótico & Ottoni, 2013; Visalberghi et al., 2017). Further, while tool use for extractive foraging allows access to otherwise inaccessible resources, these benefits are not necessarily limited to the tool user themselves. Other individuals can scrounge during tool using events if the tool-user allows it, or on resulting debris afterwards. Bearded capuchins scrounge on food remains in 24%-35% of nut-cracking events (de A. Moura & Lee, 2010; Ottoni et al., 2005), and if females routinely get access to resources in this manner such male provisioning could explain sex differences in tool use frequency. If one sex is already a less frequent tool user, for instance because of physical limitations due to size or strength, easy access to resources opened by tool use might be further incentive to use tools less frequently—thus widening the gap between sexes.

Studies of tool-using primates have, to date, rarely focused on multiple groups within the same population that differ in their patterns of tool-use, making it challenging to disentangle the morphological, social and environmental factors that may contribute to sex differences in behavior. The white-faced capuchin *(Cebus capucinus imitator)* population that inhabits the Coiba Archipelago thus provides a rare opportunity to test hypotheses about why such sex-biases arise. Despite decades of studies on white-faced capuchins across multiple field sites (Fedigan & Jack, 2012; Perry et al., 2012), stone tool use has only been documented on Coiba and Jicarón, two islands off the Pacific coast of Panamá. Here, capuchins habitually use stone tools to open sea almonds, palm fruits, hermit crabs, snails, coconuts, and other resources (Barrett et al., 2018; Monteza-Moreno, Dogandžić, et al., 2020). White-faced capuchins are generalist foragers that, across their geographic range, show a diverse repertoire of extractive foraging behaviors that are transmitted via social learning (Barrett et al., 2017; Perry, 2011). They are sexually dimorphic (♂_weight_ = 3.7 kg, ♀_weight_ = 2.5 kg; Smith & Jungers, 1997), and live in multi-male, multi-female groups with male dominance and female philopatry. Females exhibit linear dominance hierarchies, while males have no clear hierarchy beyond the alpha male (Fragaszy et al., 2004; Perry, 1998; Perry et al., 2008). In some populations, slight sex differences exist in extractive foraging behavior, with males more frequently consuming foods that require such processing (O’Malley & Fedigan, 2005) and using different techniques than females (Perry, 2009).

Hammerstone and anvil tool use by white-faced capuchins on the island of Jicarón appears to be male-biased similar to probing tool use in bearded capuchins: no instances of female tool use have been reported (Barrett et al., 2018). This is in sharp contrast to the neighboring island of Coiba, where females routinely use tools (Monteza-Moreno, Dogandžić, et al., 2020). On Jicarón, stone tool use occurs at three types of sites, classified based on debris accumulation due to activity intensity and site erasure (see Barrett et al., 2018): *i)* elusive sites with no to low accumulation, e.g, rocks in the intertidal zone where debris is washed away regularly, *ii)* small- and medium-sized accumulation sites in dry streambeds where some debris accumulates before washing away, and *iii)* high accumulation sites (from here on referred to as “anvils”), which are on higher ground away from streambeds and the shore. Anvils are used by tool-using capuchins on a near-daily basis, and are easily identifiable due to the large accumulation of tools and processed food debris. Even with extensive coastal surveys and observations going back over a decade (Ibáñez, 2011), the presence of high accumulation anvil sites appears to be limited to a small stretch of coast on Jicarón (∼1 km; Barrett et al., 2018). On Coiba, we have not found any evidence of high accumulation sites, but only of small- and medium-sized accumulation sites in streambeds.

In this study, we examine factors that could account for the apparent male bias in stone tool use in the white-faced capuchin population living on Jicarón island. We first investigate whether the observed sex bias reflects a true difference in the behavior of tool-using capuchins on Jicarón or if it is, instead, a result of biases in our sampling. Data collection on these tool-using capuchins relies heavily on camera traps placed at targeted anvil sites. This biased, targeted sampling collection might not capture female tool use for a variety of reasons, for instance, if females are outcompeted at these sites. To obtain more unbiased data in a variety of environmental conditions, we analyzed data from camera traps deployed along streambeds, where tool use occurs largely on invertebrates (e.g., snails) at ephemeral, small- to medium-sized accumulation sites, as well as in the forest interior where tool use is unlikely to occur.

Using all of our camera trap data from both Jicarón and Coiba, we then test five main hypotheses to explain the male-bias in stone tool use observed in capuchins on Jicarón (Table 1). While females are likely physically capable of tool use – juvenile capuchins are prolific tool users (Barrett et al., 2018) and females on the neighboring island of Coiba do use tools (Monteza-Moreno, Dogandžić, et al., 2020) – physical differences, including smaller body size, may reduce tool-use efficiency [**H1**] leading females to avoid this foraging strategy. Furthermore, if females, especially those carrying offspring, spend less time on the ground due to risk aversion, female tool use might occur but go undetected, either because it happens in the trees (and all camera traps are on the ground, Barrett et al., 2018) or because it occurs infrequently **[H2]**. Additionally, as anvils and hammerstones can be monopolized (Verderane et al., 2013), it is also possible that females rarely use tools because of within-group competition **[H3]**, which would be expected to be particularly high at the high accumulation anvil sites which see intensive daily use by the capuchins. Alternatively, similar to the potential male provisioning described by de Andrade Moura and colleagues (2010), female capuchins might not frequently use tools because they instead scrounge on food opened by others [**H4**]. Lastly, female white-faced capuchins might use tools on different food items than males [**H5**], which could go undetected in our initial sampling regime focused largely on one food item (sea almonds, which are the primary food item accessed by males using stone tools). Based on preliminary observations, females on Coiba appear to largely use tools to access freshwater and marine snails and palm fruits (*Bactris major* and *Astrocaryum standleyanum*), whereas sea almond processing appears more opportunistic (B. Barrett, *pers. comm.)*.

**Table 1.** Overview of hypotheses and predictions.

| *Hypotheses* | *Predictions* |
| --- | --- |
| **H1 – Morphology:** Females are physically limited in their ability to use tools | **P1a** – Female and juvenile tool use does not occur<br>**P1b** – Female and juvenile tool use is less efficient than male tool use<br>**P1c** – Females and juveniles use smaller tools than males |
| **H2 - Risk:** Females are more risk averse and spend less time on the ground than males, or do not use stone tools on the ground | **P2a** – Adult females are less frequently detected on the ground on camera traps than adult males<br>**P2b** – Adult females are less likely than males to be seen on camera traps in open spaces (i.e., streambeds and close to the coast)<br>**P2c** – Adult females carrying infants will be less likely to be seen in open spaces (e.g. streambeds and close to the coast) than females without infants |
| **H3 – Competition:** Females are not observed using tools because they are competitively excluded from (especially high accumulation) anvil sites | **P3a** – Displacements at high accumulation anvil sites are common<br>**P3b** – Females are displaced at high accumulation anvil sites<br>**P3c** – Female tool use occurs at elusive, small- and medium-sized accumulation anvils sites such as in streambeds |
| **H4 – Scrounging:** Females do not frequently use tools because they scrounge on food opened by others | **P4a** – Females scrounge on items opened by tool-users during the tool-using event<br>**P4b** – Females often forage on anvil debris left over after tool use by group mates |
| **H5 – Diet:** Females are not observed using tools because they use tools on different food items than males. | **P5a** – If we observe female tool use on Jicarón, it will (like on Coiba) be on snails in streambeds and palm fruits and not on sea almonds |

## Methods

Data used in this study come from a long-term study of the tool-using groups of white-faced capuchins living on the islands of Jicarón and Coiba, and include camera trap images collected between March 2017 and January 2023.

### Study Site

Jicarón (2 002 ha) and Coiba (50 314 ha) are two of the nine islands and 100 islets that make up Coiba National Park, a designated UNESCO World Heritage site off the Pacific coast of Veraguas Province, Panama. Both have depauperate mammalian communities compared to the mainland and lack mammalian predators (Ibáñez et al., 1997). In comparison to mainland populations, white-faced capuchins on Jicarón and Coiba live at unusually high densities (Barrett et al., 2018; Milton & Mittermeier, 1977), and are markedly more terrestrial (Monteza-Moreno, Crofoot, et al., 2020). Both Jicarón and Coiba were inhabited from 250 CE until the 16^th^ century by indigenous communities, and used as a penal colony from 1919 until 2004 (Isaza & Vrba, 2010). Currently, there is a small but constant human presence on Coiba in the form of a National Park station and police station, as well as occasional tourism. Jicarón is largely undisturbed without post-colonial human occupation.

### Study Subjects

White-faced capuchins living on Jicarón and Coiba island are unhabituated, and thus all data presented here originates from indirect means of data collection, primarily via the deployment of camera traps. The group size of the Jicarón tool-using group appears to be between 20-25 individuals, with at least 5 adult males and 5 or 6 adult females (fluctuating throughout the data collection) (Goldsborough et al., in press). Less is known about the tool-using group on Coiba island, so we cannot make estimates of group size or composition. Sex of adults and subadults can reliably be determined from camera trap footage, but estimating sex of juveniles is not reliable. Juveniles can only be sexed using clear photos of the genitalia, and this has an inherent male-bias. We identified several subadult male capuchins and juvenile male capuchins, but only one subadult female and no juvenile females, so we make no distinction between male and female subadult and juvenile capuchins in our analyses.

### Data Collection

#### Camera trapping protocol

On Jicarón, Camera traps have been placed in the tool-using group’s range continuously since March 2017 (with a gap in data collection in 2020 due to the COVID pandemic, see Supplemental Table S1 for details on camera deployments). Initially, most camera traps were placed on targeted locations at tool-using anvils and in sea almond groves. However, in 2021 we placed additional cameras along streambeds and at randomly selected locations, to reduce the bias in our sampling by capturing areas where tool use was either more ephemeral (streambeds) or entirely absent (random locations). Furthermore, in 2022 we placed a grid of 24 cameras spaced at 100 meters in the tool-using group’s range. On Coiba, we have placed camera traps in the tool-using group’s range intermittently since March 2019. We use still (Reconyx Hyperfire HC600 & HF2X) and video (Reconyx Ultrafire XR6 & XP9) camera traps, with infrared rather than white flash to minimize animal disturbance. Per trigger event, still-camera traps recorded 10 images on Rapidfire mode with ∼1 sec between images and no between-trigger delays. Video cameras recorded over a 24-hour period, and per trigger recorded one image and either a dynamic or a static video. During a dynamic video recording, the camera trap stops after 3 seconds of inactivity, and is retriggered by additional movement within 27 seconds from stopping. Dynamic videos can be of varying lengths, with a maximum length of 30 seconds, while static videos record one video of a set length of 30 seconds per trigger. Of our 15 deployed video traps, 2 were static.

On Jicarón, we surveyed 61 camera locations (11 anvil, 13 streambed, 37 random). Anvil cameras were directed at anvils (high accumulation anvil sites). Streambed cameras were placed both at suspected small- and medium-sized accumulation sites in streambeds and at other streambed locations within the tool-using group’s range without clear evidence of previous tool use. Random cameras were placed in three ways: *a)* paired with a stream camera, placed by walking approximately 15 meters away from the streambed camera perpendicular to the stream on whichever side of the stream the terrain was accessible, *b)* at random locations in the landscape within (what we estimated to be) the tool-using group’s range, or *c)* as part of a 24 camera grid placed in the tool-using group’s range at 100 meters spacing. All random cameras were deployed on an available tree facing a random direction. A total of 94 cameras (79 stills and 15 videos) were deployed during the sampling period. 46 of these camera deployments had a single sampling period, the other 15 were sampled repeatedly (between 2 and 6 deployments in the same location). Cameras were collected and deployed twice a year around March and December, and not all cameras ran until pick up so sampling was not fully continuous. Per camera, the average duration of sampling nights was 135.6 (range 10-256), totaling 12,748 sampling nights (anvil = 3,129, streambed = 3,120, random = 6,499).

On Coiba, we surveyed 9 camera locations, all of which were streambed sites where we placed still cameras for a single sampling period. In contrast to our sampling on Jicarón, streambed cameras on Coiba were placed at locations where we saw physical evidence of tool use in the streambed. Per camera, the average duration of sampling nights was 145.4 (range 28-190), totaling 1,309 sampling nights.

#### Data processing and behavioral annotation

All images and videos were compiled into sequences, where one video is a single sequence, and all bursts of images triggered within 30 seconds of one another are combined into one sequence. Sequences were coded in Agouti, an online platform for archiving and annotating camera trap data (Casaer et al., 2019).

Per sequence, we identified which animal species were visible and recorded the number of individuals per species. For white-faced capuchins, we coded the age class (adult, subadult, juvenile, infant) and sex of each individual, and assigned an individual ID whenever this was possible. We were able to successfully sex 90% of observed adult capuchins (compared to 63% of subadults and 13% of juveniles). For 25% of all observed capuchins (n_total_ = 33,465), we were unable to reliably assign sex or age, largely because these individuals were only partially in view (e.g., only a tail) or inspecting the camera so closely that they were unidentifiable. For every individual capuchin in a sequence, we coded which behavior(s) best described their activity in the majority of the sequence (for ethogram see Table S1). More active behaviors took precedent over passive behaviors, i.e., we only coded “resting” if this was truly the only thing the capuchin did in the sequence. If they foraged, even briefly, this was coded instead. We calculated the distance of each camera site on Jicarón to the coast by taking the distance from a camera’s GPS point to the nearest coastal vegetation boundary (for more details see Goldsborough et al., in press).

For analyses, we only considered sequences containing capuchins (n_Jicarón_ = 18,353, n_Coiba_ = 395). Since camera traps only capture snapshots in a limited spatial area of the environment, an absence of triggers does not necessarily mean that no capuchin was present in this area (and camera traps also do not always trigger fast enough to capture a traveling animal). We also excluded days on which cameras were deployed and collected from analyses, as the human presence on these days could affect capuchin behavior. All statistical analyses were done in R v. 4.2.2 (R Core Team, 2022).

To evaluate whether females are physically incapable of tool use or whether females and juveniles are less efficient than males (H1), we examined tool use events on Jicarón in detail through frame-by-frame coding of video observations in BORIS v. 8.14 (Friard & Gamba, 2016). For each tool-using sequence (the opening of a single item), we assessed a) the number of pounds, and b) the time in seconds required to open the item, as well as c) the number of mis-strikes. Additionally, we examined the frequency of female tool use on Coiba (where detailed frame-by-frame coding was not possible due to the distance of tool-use events from the camera).

#### Statistical analyses

To assess whether females (with and without infants) are less likely to be detected on the ground, particularly in “riskier” open areas such as streambeds or closer to the coast (H2), we employed two hierarchical logistic regressions per island, *i)* comparing the ratio of females to males and *ii)* comparing the ratio of females with infants to females without infants. Females with infants are categorized as such if they are observed providing infant care such as nursing or carrying an infant dorsally or ventrally (Table S2).

All logistic regressions were fit using Bayesian regression modelling with Stan (Stan Development Team, 2023) via the “brm” function in the brms package v. 2.16.1 (Bürkner, 2017), and included only sequences with adults present where at least one of the adults was successfully sexed (n_Jicarón_ = 6,603, n_Coiba_ = 252). First, we compared the ratio of adult females to adult males at the three location types (anvil, streambed, and random cameras) on Jicarón. In this logistic regression, the number of “successes” was the number of females observed in a sequence and the total number of adults observed in a sequence was the number of trials. This allows us to obtain the ratio of adult females to adult males. For predictors, we estimated the effect of the type of location (random, anvil or streambed, with random as reference level) and the distance of each specific location site to the coast. We also included the specific site of the camera deployment as a random effect, and compared our findings to the ratio of adult females to adult males on Coiba (where all cameras are streambed cameras).

To compare adult females with infants to those without, we fit another logistic regression with the same structure, but where the number of successes was the number of adult females with infants, and the trials the total amount of adult females in the sequence. Here, only sequences were included where at least one adult female was present (n_Jicarón_ = 2,329, n_Coiba_ = 209). We again included the specific site of the camera deployment as a random effect, as well as the month of the year to account for seasonality in births. We compared the results of this model to the ratio of adult females with infants to adult females without infants on Coiba, month was not included in the Coiba model due to limited data collection throughout the year.

To test whether females are being outcompeted at anvils (H3), we quantified competition at anvils on Jicarón through the number of displacements observed, where a displacement means that one individual supplants another from the tool anvil and/or hammerstone (see Table S1). We considered the actual number of displacements in relation to the opportunity for displacement, quantified as the number of sequences when individuals of the age-sex class were at an anvil with at least one other individual present. We further considered the opportunity for adult females to use tools by examining how frequently adult females were observed at anvil sites without adult males present (under the assumption that this would be a situation of low competition).

To examine whether female**s** do not frequently use tools because they scrounge on food opened by others (H4), we compared the likelihood of different age-sex classes to scrounge during tool-using events or on anvil debris by comparing the frequency of observed scrounging to the opportunity to scrounge. For scrounging during tool-using events, the number of opportunities to scrounge is reflected by the number of sequences a capuchin was present at an anvil together with a tool-user. Opportunity to forage on anvil debris were all sequences where a specific age-sex class was present at the anvil, regardless of how many other capuchins were present.

In a first investigation of whether females use tools on different food items than males (H5), we used data from Coiba as a comparator. We deployed camera traps in the streambeds on both Jicarón and Coiba to quantify tool use away from anvils (i.e., in small- to medium deposition sites. Due to the limitations of camera traps in inferring diet (e.g., not capturing all items consumed, bias towards observing consumption of larger food items), our examination of sex differences in diet is necessarily limited.

All models were fit with regularizing Normal(0,1) priors for intercepts and predictors, and exponential(1) priors for standard deviations of varying effects. We performed a prior predictive simulation to visualize the predictive implications of the priors and evaluate identifiability of parameters. We ran the final models with 3 chains, each having 3,000 iterations, including a warm-up period of 1500 iterations per chain. Our models were stable with large effective sample sizes (Bulk_ESS and Tail_ESS over 1,000 for all estimates; Bürkner, 2017) and Rhat values < 1.01 (Vehtari et al., 2021). For all models, Pareto k estimates were below 0.5. We used the posterior predictive check function to visually assess model fit and confirm our choice of priors. For full model specifications and details see the Supplementary Materials and associated reproducible R code (https://doi.org/10.5281/zenodo.831626).

## Results

### H1: Females are physically limited to use tools

We observed no instances of female tool use on Jicarón in the 5 years of camera trap data we have collected (12,748 sampling days, 18,353 sequences with capuchins), compared to 1453 tool use events by adult males, 694 by subadults and 2594 by juveniles. On Coiba, we observed adult females use tools 11 times (1,309 sampling days, 395 sequences with capuchins), which is more than we observed for adult males (3 times), subadults (4 times), or juveniles (3 times).

Based on a preliminary comparison of the time and number of pounds required to successfully open an item in 339 tool use sequences on Jicarón, juveniles are less efficient (mean from raw data 5.9 pounds [range 1-34], 24.8 seconds [range 3.1-180.8]) than adult males (mean 3.8 pounds [1-14], 10.2 seconds [2.6-44.8]). Nonetheless, juveniles used the same hammerstones as adult males (mean weight of all measured hammerstones on Jicarón is 660g [n = 174]), and longer processing times by juveniles can also be a result of less experience using tools.

### H2: Female tool use is limited due to increased risk aversion

#### Ratio of adult females to adult males

Adult female capuchins were equally as likely to be seen on the ground as adult males at random cameras. Our model comparing the ratio of adult females to adult males estimated considerable differences between all three location types (Figure 1, see Table S3 for estimates). Based on our estimates of the true ratio of adult females to adult males in the group (5 adult males and 5 or 6 adult females; 0.50-0.55), both sexes were equally as likely to be seen on the ground at random cameras (Median = 0.56 [95% CI 0.50, 0.62]). The ratio of adult females to adult males at streambed cameras (Median = 0.38 [95% CI 0.30, 0.47]) was considerably lower than at random cameras (posterior probability exceeding 95%). At anvils we observed the lowest ratio of adult females to adult males (Median = 0.33 [95% CI 0.21, 0.47]), considerably lower than at random cameras, but comparable to streambed cameras. On Coiba, where all cameras were placed on streambeds, the ratio of adult females to adult males was higher than at any of the location types on Jicarón (Median = 0.76, 95% CI [0.61-0.85], Table S4 and Figure S2).

**Figure 1.**
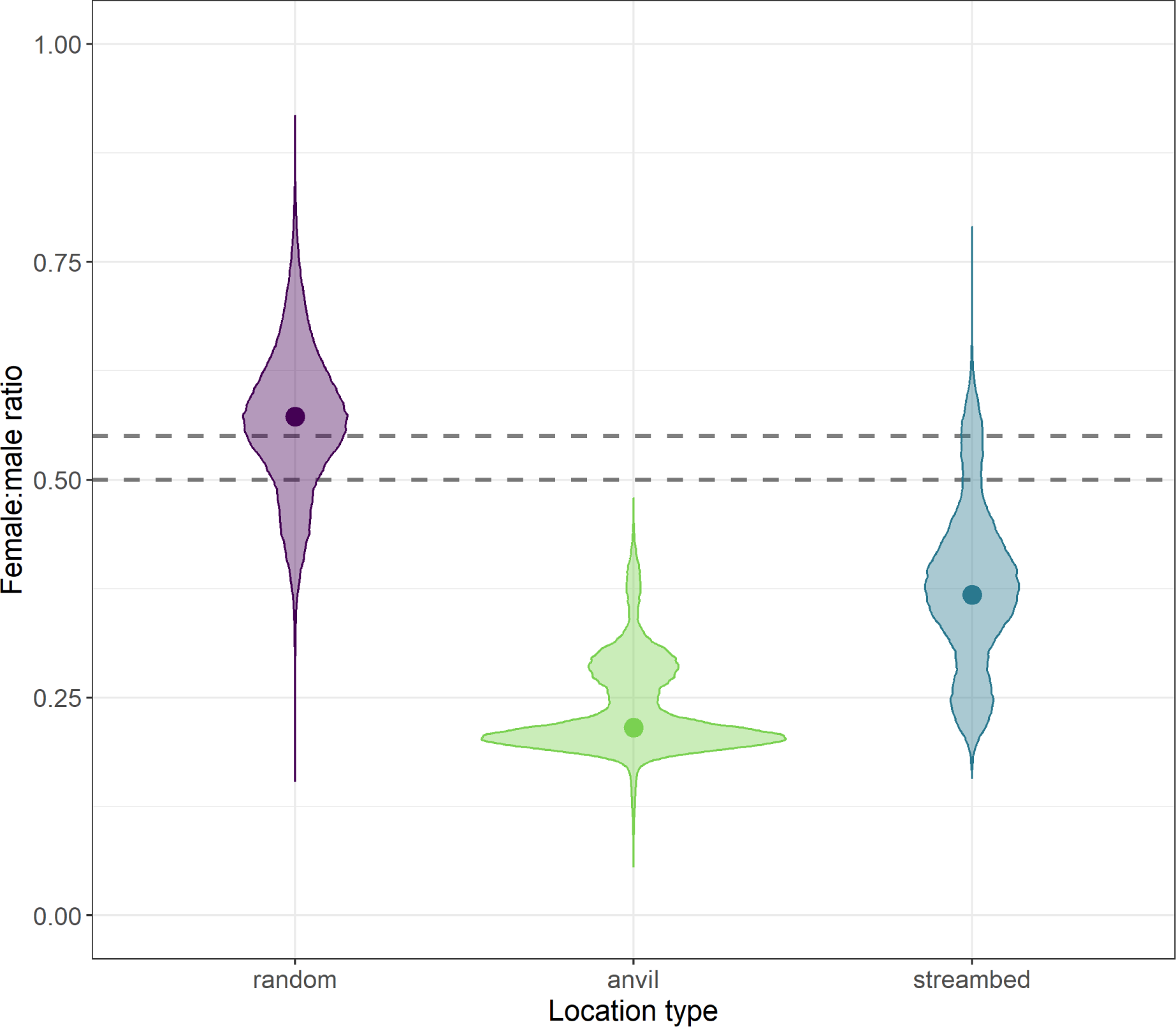
Model estimates and mean values of the ratio of adult females to adult males at each location type (random, anvil, streambed). Model estimates are reflected by violin plots, and observed means from data are represented as points. Dashed horizontal lines represent range of the estimated true ratio of adult females to adult males based on how many distinct individuals are present in the group (5 or 6 adult females and 5 adult males).

The interaction effect of distance to coast and anvil sites (Median =1.00, 95% CI [−0.17, 2.12]) has a 96% probability of being positive (> 0) and is a moderate effect, indicating that adult females were slightly less likely to be seen at anvil sites closer to the coast compared to those further inland (Figure 2). At both random (Median =0.05, 95% CI [−0.07, 0.16]) and streambed cameras (Median =0.23, 95% CI [−0.22, 0.67]), the interaction with distance to coast was also estimated to be positive (77% posterior mass for random cameras, 84% for streambed cameras), but these positive relationships were weaker and less reliable. Note that anvil sites are limited to the coast, and the range of camera distances varied between the location types (random = 8.1-328.3 m, anvil = 0.8-30.8 m, streambed = 25.2-140.4 m).

**Figure 2.**
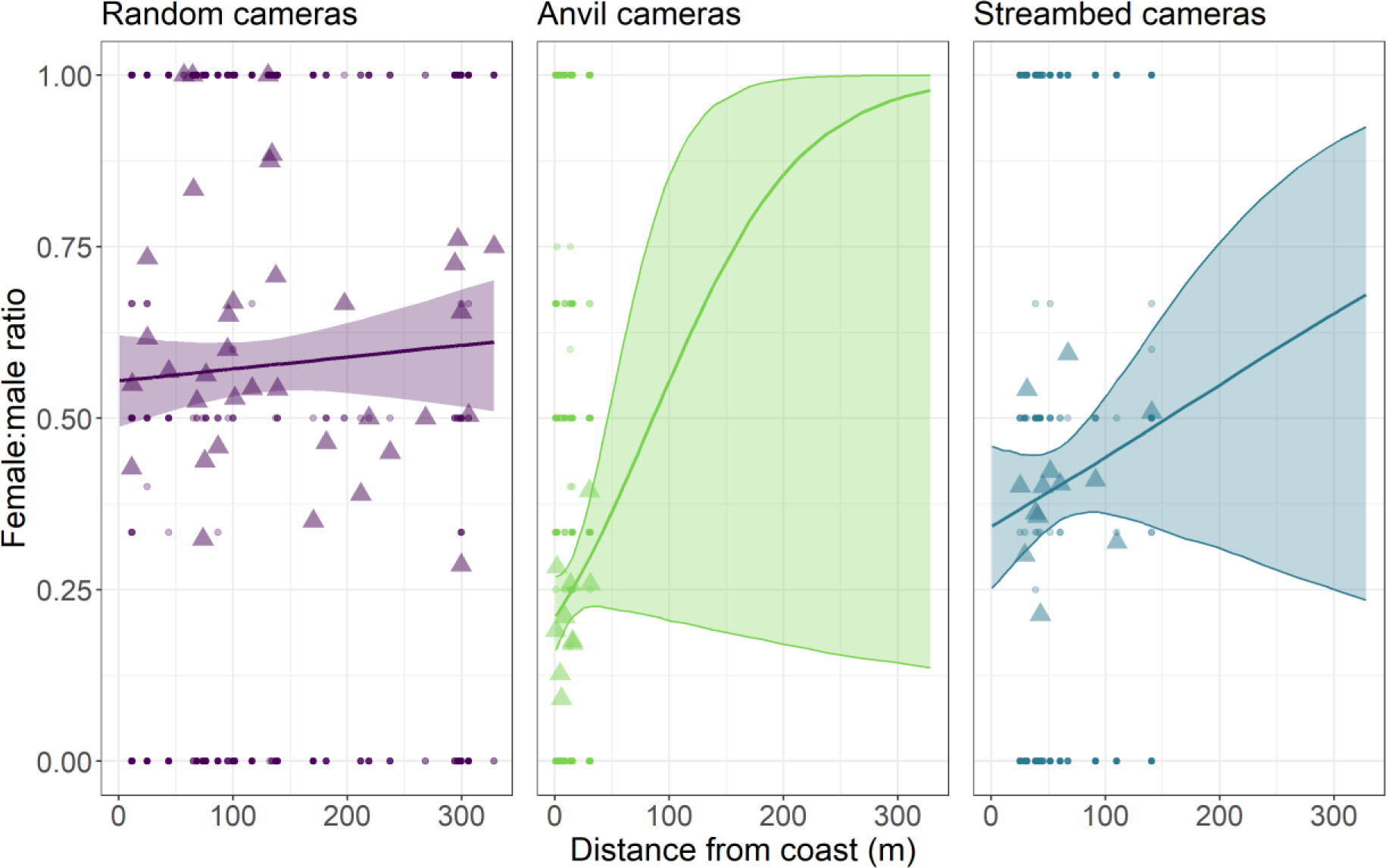
Marginal effects of the type of camera location (random, anvil, or streambed) and the distance to coast on the female to male ratio. Lines indicate the marginalized median sex ratio from model predictions, with the ribbon reflecting the 95% CI. Triangles represent the empirical means per camera trap. Circles are raw datapoints.

#### Ratio of adult females with infants to adult females without infants

There were no strong differences in the ratio of adult females with infants to the ratio of adult females without infants between any of the three location types (Figure 3, see Table S5 for model estimates). Adult females with infants were less likely to be seen than adult females without infants at random cameras (Median = 0.21, 95% CI [0.15, 0.28]), streambed cameras (Median = 0.27, 95% CI [0.18, 0.38]) and anvils (Median = 0.16, 95% CI [0.08, 0.29]). We do not know the true ratio of adult females with infants to adult females without infants, so can only compare between locations. At the streambed cameras on Coiba, the ratio of adult females with infants to adult females without infants was lower than at any of the location types on Jicarón (Median = 0.08, 95% CI [0.01, 0.37], Table S6 and Figure S5).

**Figure 3.**
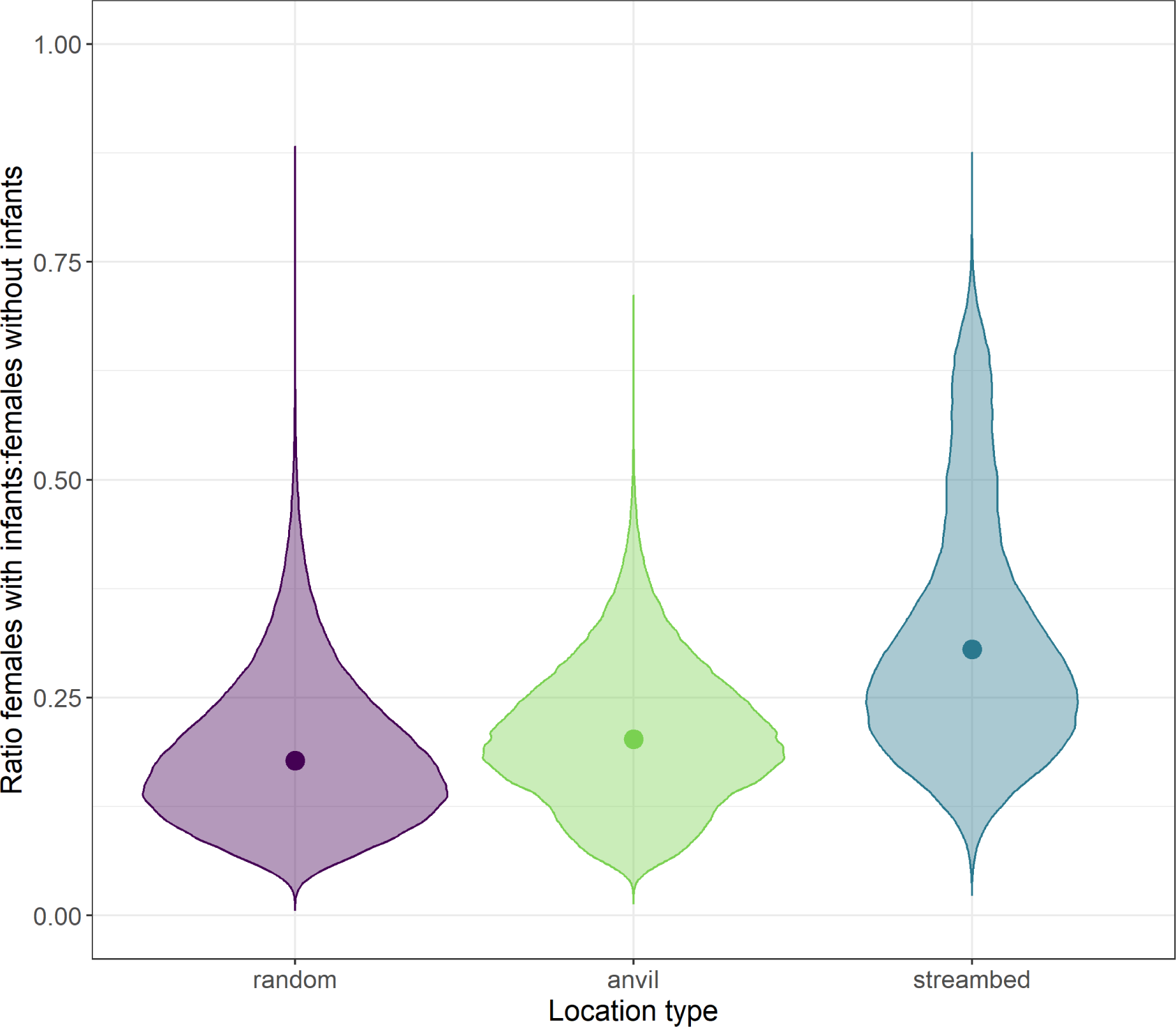
Model estimates and mean values of the ratio of adult females with infants to adult females without infants at each location type. Model estimates are reflected by violin plots, and observed means from the data are represented as points.

We find that adult females with infants were more likely to be seen at anvil sites closer to the coast than at those further inland (Figure 4). The interaction effect of distance to coast and anvil sites (Median =-0.52, 95% CI [−1.89, 0.97], 75% of posterior <0) is a moderate negative effect. While at random cameras, the negative effect is smaller and less reliable (Median =-0.08, 95% CI [−0.25, 0.09], 80% of posterior <0). At streambed cameras, the positive relationship is reliable and of moderate size (Median = 0.75, 95% CI [0.12, 1.36], 99% of posterior >0). Thus, adult females with infants were considerably less likely to be seen at streambed cameras closer to the coast than at streambed cameras further inland.

**Figure 4.**
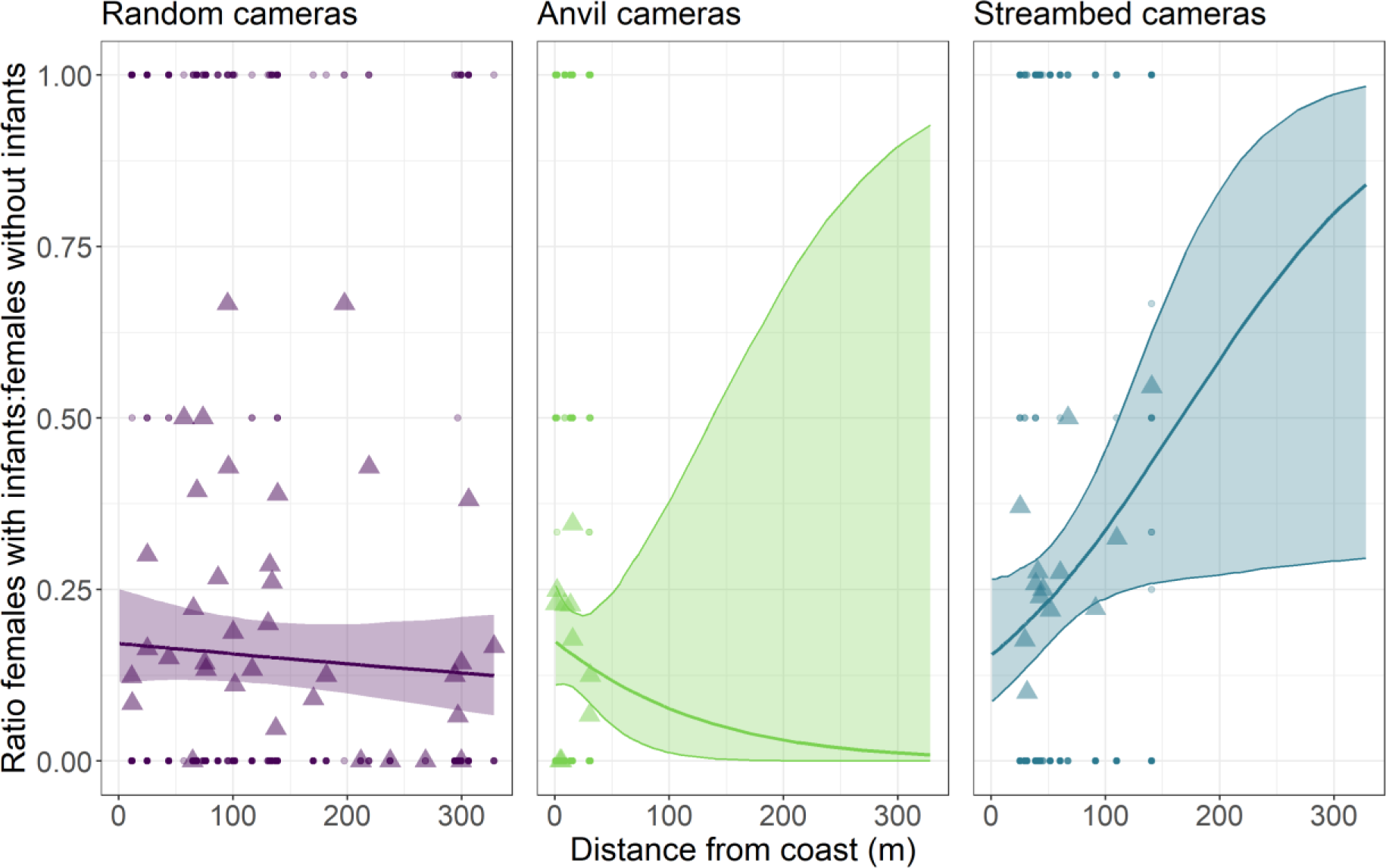
Marginal effects of the type of camera location (random, anvil, or streambed) and the distance to coast on the ratio of adult females with infants to females without infants. Lines indicate the marginalized median sex ratio from model predictions, with the ribbon reflecting the 95% CI. Triangle points represent empirical means per camera trap, and round points are raw datapoints.

Both models indicated considerable variation between camera locations and the second model indicated monthly variation in the ratio of females with infants to females without females (see Supplementary Figures S1 and S3 for model estimates per camera location, Figure S4 for model estimates per month).

### H3: Females are outcompeted at tool-use anvils

Direct competition at anvils on Jicarón appears to be extremely low. We observed capuchins at anvil sites in 12,274 sequences, and in less than half of those (5,184 sequences) more than one individual was present, meaning there was an opportunity for displacements to occur. Despite these opportunities, displacements were rare, occurring only 92 times (1.77% of all displacement opportunities).

Direct competitive dynamics over anvils likely cannot explain why adult females do not engage in stone-tool use. They were rarely displaced themselves (Figure 5), and were also observed to displace other (primarily juvenile) members of the group. Further, adult females were observed at anvil cameras in 736 sequences without any adult males present who could outcompete them (under the assumption that females would mostly be outcompeted by adult males and not by juveniles). In 638 of these sequences there were also no juveniles using tools, so the female(s) could easily access the anvil. On Coiba, we observed no displacements, despite 141 opportunities for displacement (i.e., more than one capuchin present in a sequence).

**Figure 5.**
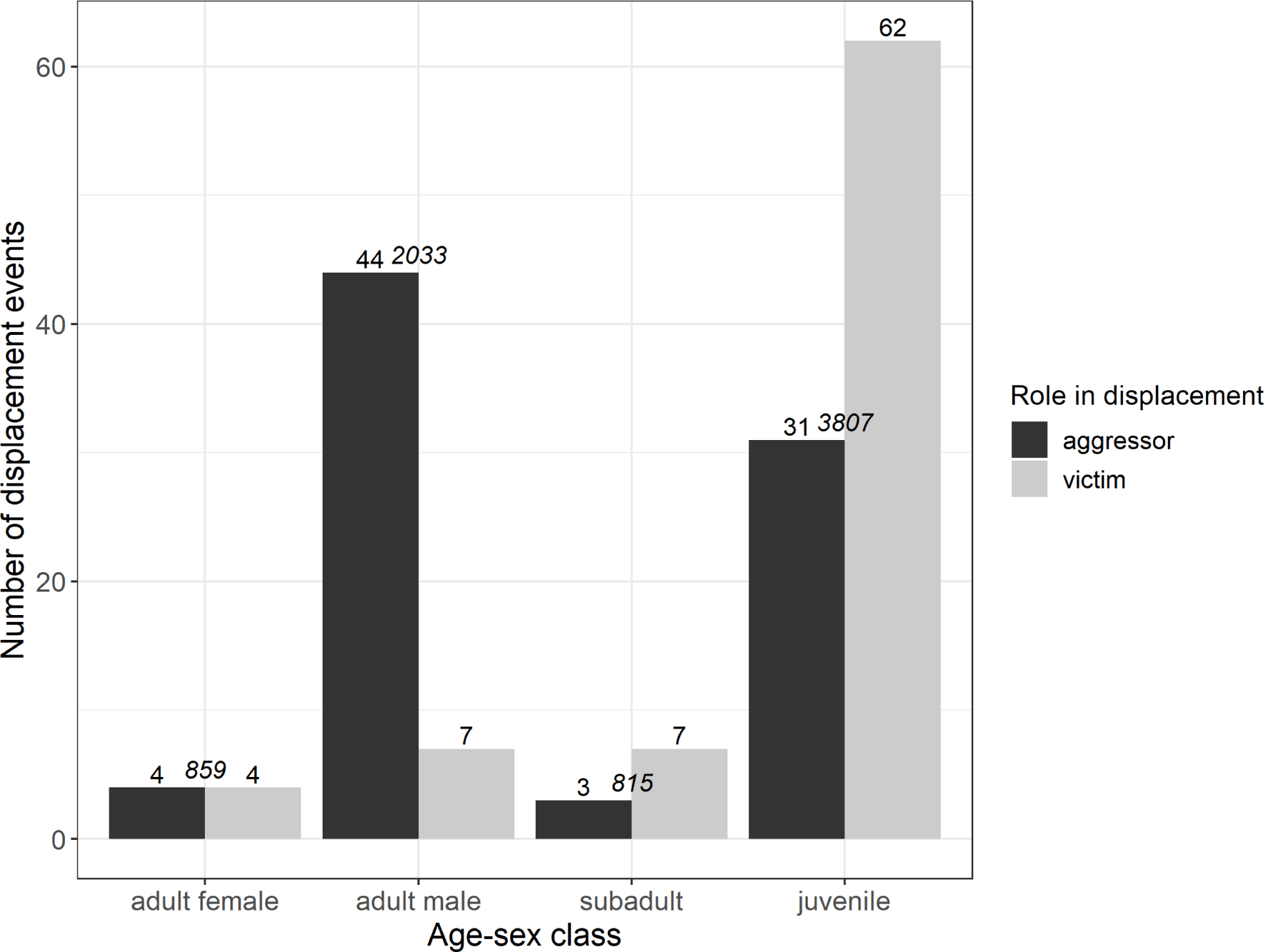
Number of times each age-sex class was observed being the aggressor (black bar) or victim (grey bar) in a displacement event. Events when we could not identify the age-sex class of the aggressor (n = 10) or victim (n = 12) are excluded. Italicized numbers reflect the number of events where at least two individuals were present in a sequence, representing opportunities for displacement events to have occurred.

### H4: Females frequently scrounge on food opened by others

Resource scrounging during tool-using events was rarely observed, occurring in 3.5% of all opportunities (Table 2), and adult females were only observed scrounging on two occasions (1% of their opportunities). The lack of scrounging does not appear to be related to tool-users refusing to tolerate scrounging. Foraging on anvil debris left over after tool use was similarly rare (occurring in 5.1% of opportunities). In both situations, juveniles, subadults, and adult males were more likely to scrounge during tool use or forage on anvil debris than adult females.

**Table 2.** Proportion of times each age-sex class was observed scrounging during a tool-using event or on anvil debris afterwards.

| *Scrounging during tool use events* |  |  |  |
| --- | --- | --- | --- |
| Age-sex class | Observed scrounging events | Scrounging opportunities | Percentage of opportunities spent scrounging |
| Adult female | 2 | 205 | 1% |
| Adult male | 17 | 801 | 2.1% |
| Subadult | 4 | 348 | 1.1% |
| Juvenile | 83 | 1396 | 5.9% |
| Unknown | 5 | 391 | 1.3% |
| *Scrounging on anvil debris* |  |  |  |
| Adult female | 43 | 1224 | 3.5% |
| Adult male | 211 | 3900 | 5.4% |
| Subadult | 71 | 1658 | 4.3% |
| Juvenile | 761 | 9954 | 7.6% |
| Unknown | 74 | 6001 | 1.2% |

### H5: Females use tools on different resources than males

Despite a total of 3,120 sampling nights at 13 streambed camera locations, we did not observe any tool use on snails or other bivalves by adult females on Jicarón. In contrast, all observed female tool use events (n= 11) by females on Coiba were on snails or other invertebrates in the streambed. Males on Jicarón were mostly observed using tools at high accumulation anvil sites, which are away from streambeds and close to sea almond groves (1,439 instances of tool use across 3,129 trapping days). At streambeds, adult males also used tools to process invertebrates, but this was less frequently observed (12 instances of tool use across 3,120 trapping days).

## Discussion

Stone-tool use by white-faced capuchins on Jicarón island is strongly male-biased. We observed no instances of female tool use on Jicarón in more than 5 years of data collection with camera traps at different locations in the tool-using group’s range (totaling 12,748 camera trapping days), and despite the fact that females regularly use stone tools on the neighboring island of Coiba. Differences in physical capacity, risk-aversion, competition, or scrounging behavior cannot account for the observed differences in tool use among males and females in this population.

Sampling biases are unlikely to explain the observed differences in female vs. male tool-use behavior, given the temporal and spatial depth of our data. Female capuchins are less frequently seen at anvil sites than male capuchins, but at the 13 cameras placed at streambeds and 37 cameras placed on random locations we saw comparable female and male activity. On the neighboring island of Coiba, capuchins—including adult females—use tools at small- to medium-sized accumulation sites in streambeds. Although we only monitored a subset of the streambeds on Jicarón, we would have captured evidence of female tool use if female capuchins on Jicaron use tools at comparable rates to males or females living on Coiba. Our findings indicate that the sex bias in tool use we observe in our data almost certainly reflects a real behavioral difference, and that adult female capuchins in the tool-using group on Jicarón rarely use stone tools (if at all).

### Adult females are physically capable of tool use

Adult female capuchins are almost certainly physically capable of stone tool use, as evidenced by the frequent tool use of juveniles who, although smaller than adult females, regularly use the same hammerstones as adult males, and tool use by adult females on the neighboring island Coiba. Comparable to in macaques (Gumert et al., 2011) and *Sapajus* capuchins (Spagnoletti et al., 2011), adult female tool-users might employ different size hammerstones and/or be less efficient than adult males due to differences in size or strength. In other populations of white-faced capuchins, adult males engage in more extractive foraging of *Luehea* and *Sloana* fruits than adult females, perhaps because of the strength required to break fruits off their stems (Perry, 2009). However, as we observed juveniles successfully using the same size hammerstones as adult males, physical strength is unlikely to be a limitation for tool use by adult females in this population.

### Adult females have opportunity to use tools

Adult females, both with and without dependent offspring, were just as likely to be seen on the ground as adult males at random cameras in the vegetation, similar to findings by A. de Moura and colleagues (2010). The lower likelihood to observe adult females at anvil sites and streambeds is likely not due to risk, but rather a consequence of the lack of tool use by females and/or less interest in consuming sea almonds than males. Both sexes of capuchins on Jicarón appear to have low risk aversion, given their generally high degree of terrestriality and investigation of camera traps (Monteza-Moreno, Crofoot, et al., 2020), which could be related to the absence of mammalian predators on the island (Ibáñez et al., 1997). However, even though males and females are similarly terrestrial, females could still show different behavior while on the ground (e.g., be more vigilant and less likely to engage in conspicuous behavior like tool use) due to a higher perceived risk than males. Our finding that adult females are less likely to be seen at cameras closer to the coast compared to further inland, particularly at anvil sites, also appears unrelated to risk: sea almonds, the main food type stone tools are used to access on Jicarón, are a coastal species that is abundant along the coast and absent inland. Furthermore, the ratio of adult females with infants to adult females without infants was comparable between location types. Adult females with infants being more common at anvils closer to the coast could be due to the social dynamics in the group: adult males spend more time using tools at coastal anvils than further inland, since sea almonds are a coastal species, consequently the whole group might spend more time in this area than at less suitable anvils. The opposite relationship we observe for streambed cameras, females with infants being more likely to be seen inland, could be capturing that some feature of the streambeds is mitigated by distance to the coast (for instance, freshwater shrimp are more common upstream than at the entrance of streams where water is more brackish).

Adult females were less likely to be seen at anvil sites than at streambeds and random cameras, which would suggest that potentially they are being outcompeted at anvil sites. However, displacements at anvils were strikingly uncommon. In nearly half of all tool-using events, only one tool-using individual was seen in the camera trap photos. This suggest that either competition at anvils is not particularly high (in contrast to Verderane et al., 2013), or competition is resolved via behavioral mechanisms other than displacement such as increasing inter-individual foraging distances. The apparently solitary nature of stone tool use makes it unlikely that it serves a display function driven by sexual selection as suggested in bearded capuchins (de A. Moura & Lee, 2010). Additionally, adult females evidently have access to anvil sites, which is reflected by the large number of sequences with adult females being the only capuchin in frame at an anvil site.

### Adult females might have different diets than adult males

The near absence of scrounging and consumption of anvil debris suggests that adult females may spend less time at anvils because they are not interested in consuming the resources available there (unlike findings in *Sapajus* capuchins, de A. Moura & Lee, 2010; Ottoni et al., 2005). At high accumulation anvil sites on Jicarón, sea almonds are the primary resource consumed. On Coiba, female tool use is largely focused on snails, palm fruits and other invertebrates in streambeds—sea almond consumption is opportunistic and, unlike on Jicarón, does not involve active transport of food items, but instead focuses on washed up fruits. Further research, for instance by using DNA barcoding, is required to test whether female and male capuchins on Jicarón truly have different diets, as the data we currently have available do not allow us to conduct detailed dietary comparisons. Previous research on other populations of white-faced capuchins did find sex differences in diet and foraging, with females spending more time foraging than males, females consuming more embedded invertebrates than males, and pregnant and lactating females focusing on foods that required little handling (Rose, 1994). Later research indicated that apparent sex differences in foraging, specifically differences in invertebrate foraging, might be a consequence of sex related variation in color vision: male capuchins all have dichromatic color vision, while females can have di- or trichromatic vision (Melin et al., 2010). Trichromatic females spent more time foraging on insects because they can spot them less well than males and dichromatic females, who showed similar foraging times but were more efficient foragers. However, this difference may be mediated by energetic requirements of reproduction, as a later study found that nursing females foraged less overall than cycling females and di- and trichromats did not differ in time spent foraging on different food types (DePasquale et al., 2021).

### Development of male-biased stone tool use

Our results indicate that adult female capuchins on Jicarón are physically capable of tool use, frequently use terrestrial substrates where tool-use would be possible, and have access to anvils where such hammerstone tool-use occurs. Nonetheless, we find no evidence that females in this population use tools. Differences in diet between sexes might lead to female tool use being undetected, but even if females use tools on snails instead of sea almonds female tool use would likely have been detected by our camera traps in streambeds. However, the intertidal zone is also an area where tool use occurs on Jicarón (*pers. obs*) where we are unable to record behavior with camera traps, so if female tool use would occur exclusively on marine resources in the intertidal we would miss it with our current sampling regime. Yet, we would expect tool use to not be limited to one area, but used flexibly in different locations like streambeds or at high accumulation anvil sites.

Given our current data, it appears most likely that adult females on Jicarón do not use tools at all, or only very rarely, comparable to the sex bias in bearded capuchin’ probe tool use (Falótico et al., 2021). Similar to the probing tool use, female capuchins on Jicarón might engage with tools as juveniles yet stop or reduce the frequency of this behavior once they mature. As we cannot reliably identify female juveniles, this hypothesis is as of yet untestable in this study system, but seems likely given the large number of juveniles we observe using tools. The mechanism triggering a cessation in tool use by mature female capuchins could be, as shown in the bearded capuchins, a motivational sex bias (Falótico et al., 2021), potentially driven by a higher tendency of male capuchins to engage in pounding behavior (Perry, 2009). Alternatively, the onset of the sex difference in tool use may be linked to females caring for dependent offspring. Carrying an infant might interfere with an individual’s ability to use stone tools due to limitations to movement and increased risk of the infant of falling off or being injured. However, in bearded capuchins, females are seen nut-cracking while carrying offspring (Mangalam et al., 2018), and we have some observations of male capuchins using tools while carrying a juvenile dorsally or even ventrally (*pers. obs*), suggesting infant carrying and stone tool use are not wholly incompatible.

### Possible explanations for strong differences in sex-biased tool use between islands

Why female capuchins on Jicarón do not use tools, and both sexes on Coiba use tools remains unclear. Whether this stark behavioral difference is a flexible behavioral adaptation to different ecological conditions (i.e. dietary richness, interspecific competition, population density) between islands, or a side effect of some other difference is social behavior or genetic difference remains unclear. One possible explanation lies in a reversal or change of dispersal tendency, as suggested by Barrett and colleagues (2018). Changes from a species’ “common” dispersal tendency – male-biased dispersal in white-faced capuchins (Jack & Fedigan, 2004) – to something else, has been observed in other populations of white-faced capuchins (Jack & Fedigan, 2009) and can occur under high density conditions (Clutton-Brock & Lukas, 2012)—conditions which appear met on Jicarón island. However, it is challenging to make predictions as to exactly when dispersal patterns can switch as there remains a gap linking quantitative theory of dispersal and empirical study of sex-biased dispersal (Li & Kokko, 2019). Important factors likely affecting sex-biased dispersal include population density, competition for limited resources, timing of breeding, dispersal mortality, and resource-sharing/territorial overlap, and inbreeding avoidance or attraction: all things that might differ in island, compared to well-studied mainland, populations. If females learn how to use tools and do so rarely, but subsequently disperse from their natal tool-using group, then tool use might be less likely to spread. Alternatively, differential sex-linked mortality could explain the observed pattern on Jicarón. Rare, deleterious traits are common on islands, including Coiba, where albinism has been reported in white-faced capuchins (Duquette et al., 2015).

## Conclusion

Tool use by adult female capuchins in the group of stone tool using capuchins on Jicarón appears to be entirely absent, rather than undetected. None of the conventional hypotheses can explain the lack of female tool use: females are physically capable of tool use, present at tool-using sites, and have access to tool-using materials and resources. Despite having ample opportunity, females show also little interest in scrounging on leftovers of stone tool use. Although we are limited in our understanding of the development of this sex bias since we cannot yet reliably sex juveniles, our findings suggest that adult female white-faced capuchins on Jicarón might have a different diet than male capuchins, or learn how to use tools as juveniles but are limited by infant-care as adults, migrate out of the tool-using group or experience differential sex-linked mortality. Further evidence, particularly DNA analyses of diet and relatedness, is required to illuminate which of these hypotheses might hold true. The mechanisms causing these sex differences likely have different implications for if and how tool-use is culturally transmitted between groups. Our findings show that there is great intragroup variation in the tool use behavior on Jicarón island, which might explain why there is also such stark intergroup variation.

## Ethics Statement

Data for this study was collected as minimally invasively as possible. We obtained ethical permission for this study from the relevant authorities, namely STRI and the Ministerio de Ambiente, Panama (scientific permit no. SC/A-23-17, SE/A-98-19, and corresponding renewals and addenda).

## Author Contributions

ZG: Conceptualization, Methodology, Validation, Formal analysis, Investigation, Data Curation, Writing – Original Draft, Visualization. MCC: Conceptualization, Writing— Review & Editing, Supervision, Project Administration, Funding Acquisition. BJB: Conceptualization, Methodology, Investigation, Resources, Data Curation, Writing—Review & Editing, Supervision, Project administration, Funding acquisition.

## Acknowledgments

We want to thank the staff at STRI, particularly the field station on Rancheria, for their support and assistance, and Lucia Torrez, Katrin Dieter and Angie Ruiz for supporting the logistics of this research. Additionally, Evelyn Del Rosario, Eliezer Vega, Pedro Luis Castillo, Zarluis Miguel Mijango, Lilisbeth Rodríguez, Tobias Begeer, Chris Dillis, Tamara Dogandzic, James Tejedor Chaves, and Juan Rojas Garrido participated in fieldwork. We also wish to thank Claudio Monteza-Moreno for his invaluable assistance with the data collection of this study, administration of the project, and for providing feedback on the manuscript. We are grateful to Alison Ashbury for writing suggestions and feedback, and to Meredith Carlson for providing data on hammerstone weights.

