## Supplemental Information for "Male-biased stone tool use by wild white-faced capuchins (*Cebus capucinus imitator*)"

**Table S1.** Overview of camera deployments

| **Location** | ***R1*** *Mar-Jul 2017* | ***R2*** *Jul-Dec 2017* | ***R3*** *Dec 2017-Mar 2018* | ***R4*** *Mar-Jul 2018* | ***R5*** *Jul-Dec 2018* | ***R6*** *Dec 2018-Mar 2019* | ***R7*** *Mar-Jul 2019* | ***R8*** *Jul-Dec 2019* | ***R10*** *Jul-Dec 2021* | ***R11*** *Dec 2021-Jul 2022* | ***R12*** *May 2022-Jan 2023* |
|---|---|---|---|---|---|---|---|---|---|---|---|
| ***Anvil cameras*** | | | | | | | | | | | |
| CEBUS-01 | V | S | V | | V | V | | | | | |
| CEBUS-02 | S | V | S | S | S | S | | | | | |
| CEBUS-04 | | | | S | S | S | | S | | | |
| CEBUS-05 | | | S | | S | S | | S | | | |
| CEBUS-06 | | | | V | V | V | | | | | |
| CEBUS-07 | | | | | | V | S | | | | |
| CEBUS-08 | | V | V | V | V | V | | | | | |
| CEBUS-09 | | S | S | S | S | S | | | | | |
| CEBUS-10 | | | | | | | S | | | | |
| SURVEY-CEBUS-07-03 | | | V | | | | | | | | |
| SURVEY-CEBUS-17-03 | | | | S | | | | | | | |
| ***Streambed cameras*** | | | | | | | | | | | |
| J-STREAM-01 | | | | | | | | | S | S | |
| J-STREAM-02 | | | | | | | | | S | S | |
| J-STREAM-03 | | | | | | | | | S | S | |
| J-STREAM-04 | | | | | | | | | S | S | |
| J-STREAM-05 | | | | | | | | | S | S | |
| J-STREAM-06 | | | | | | | | | S | S | |
| J-STREAM-07 | | | | | | | | | | S | |
| J-STREAM-08 | | | | | | | | | | S | |
| J-STREAM-09 | | | | | | | | | | S | |
| J-STREAM-10 | | | | | | | | | | S | |
| J-STREAM-11 | | | | | | | | | | S | |
| J-STREAM-12* | | | | | | | | | | | |
| JIC-STREAM-DISC-T-1 | | | | | | S | | | | | |
| SURVEY-CEBUS-15-04 | | | | S | S | | | | | | |
| ***Random cameras*** | | | | | | | | | | | |
| *Non-grid* | | | | | | | | | | | |
| J-RAN-01 | | | | | | | | | | S | |
| J-RAN-02 | | | | | | | | | | S | |
| J-RAN-03 | | | | | | | | | | S | |
| J-RAN-04 | | | | | | | | | | S | |
| J-RAN-05 | | | | | | | | | | S | |
| J-RAN-06* | | | | | | | | | | S | |
| J-RAN-07 | | | | | | | | | | S | |
| J-RAN-08 | | | | | | | | | | S | |
| J-RAN-09 | | | | | | | | | | S | |
| J-RAN-10 | | | | | | | | | | S | |
| J-RAN-11 | | | | | | | | | | S | |

| | | | | | | | | |
|---|---|---|---|---|---|---|---|---|
| J-RAN-12 | | | | | | | S | |
| SURVEY-CEBUS-08-01 | S | | | | | | | |
| SURVEY-CEBUS-16-01 | | S | | | | | | |
| *Grid* | | | | | | | | |
| TU-135 | | | | | | | | S |
| TU-137 | | | | | | | | S |
| TU-141 | | | | | | | | S |
| TU-151 | | | | | | | | S |
| TU-153 | | | | | | | | S |
| TU-154 | | | | | | | | S |
| TU-155 | | | | | | | | S |
| TU-156 | | | | | | | | S |
| TU-157 | | | | | | | | S |
| TU-158 | | | | | | | | S |
| TU-166 | | | | | | | | S |
| TU-169 | | | | | | | | S |
| TU-170 | | | | | | | | S |
| TU-171 | | | | | | | | S |
| TU-172 | | | | | | | | S |
| TU-173 | | | | | | | | S |
| TU-182 | | | | | | | | S |
| TU-183 | | | | | | | | S |
| TU-184 | | | | | | | | S |
| TU-185 | | | | | | | | S |
| TU-186 | | | | | | | | S |
| TU-187 | | | | | | | | S |
| TU-188 | | | | | | | | S |
| TU-190 | | | | | | | | S |

* Indicates a camera that was deployed but lost or destroyed

**Table S2.** Ethogram of relevant behaviors (part of larger ethogram).

| Behavior | Description |
|---|---|
| BS: Aggression | Aggressive towards another capuchin (or species). Fight, bare teeth threat, stacked coalitionary aggression, open mouth threat. This includes when an individual supplants another (specified in comment if supplanting from anvil, tool, or food). |
| BS: Infant Care | Nursing or caring for infant, including dorsal or ventral carrying of an infant. |
| BS: Submissive | Being fearful or submissive toward an aggressive individual (also code when being supplanted by another) |
| BS: Visually foraging | Looking around leaves/sticks/stream/leaf litter for food, without any physical interaction |
| BS: Tolerated scrounging | Individual takes food to eat from another individual (both tool use and regular foraging) and is tolerated. Specified in comment whether food sharing appears intention (i.e., handing over food) |
| F: Almendra flesh | Consuming almendra exocarp, no tools used or endocarp eaten |
| F: Anvil debris | Foraging opened food at anvil when tool user not there (also applies to other species) |
| F: Coco flesh | Consuming coconut meat (white part), no tools (but smashing the coconut on an anvil is part of this). |
| F: Coco water | Drinking coconut water, typically smaller coconuts. |
| F: Fruit | Foraging fruit (comment if known) |

| | |
|---|---|
| F: Insect | Foraging insects (searching leaf litter using hands, chewing sticks for embedded insects, straight up eating an insect). |
| F: Other | Foraging other (comment if known) |
| F: Unknown | Foraging unknown object |
| TAF: Almendra | Foraging w/ stone tool: *Terminalia catappa*/sea almond |
| TAF: Coconut | Foraging w/ stone tool: coconut |
| TAF: Embedded insect | Foraging w/ stone tool: embedded insect in stick |
| TAF: Halloween crab | Foraging w/ stone tool: Halloween crab |
| TAF: Hermit crab | Foraging w/ stone tool: hermit crab |
| TAF: Other | Foraging w/ stone tool: other (comment if known) |
| TAF: Palm fruit | Foraging w/ stone tool: palm fruit |
| TAF: Snail | Foraging w/ stone tool: snail |
| TAF: Unknown | Foraging w/ stone tool: unknown object |

BS indicates Behavioral State, F indicates Foraging without tools and TAF foraging with stone tools

**Model 1: Comparing adult female:adult male ratio between location types on Jicarón**

| | ***Estimate*** | ***CI_95_low*** | ***CI_95_high*** |
|---|---|---|---|
| *Group-Level Effects* | | | |
| Camera location | 0.37 | 0.25 | 0.53 |
| *Population-Level Effects* | | | |
| Intercept | 0.26 | 0.02 | 0.50 |
| Location type: Anvil | -0.97 | -1.60 | -0.38 |
| Location type: Streambed | -0.75 | -1.11 | -0.39 |
| Distance to coast | 0.05 | -0.07 | 0.17 |
| Interaction Anvil x Distcoast | 0.96 | -0.18 | 2.13 |
| Interaction Streambed x Distcoast | 0.23 | -0.24 | 0.66 |

**Table S3.** Posterior mean model estimates of model comparing sex ratio across location types on Jicarón, a Bayesian logistic mixed model. Model's explanatory power is moderate ($R^2$ = 0.20, 95% CI [0.19, 0.22]). All estimated effects are on logit scale. Random cameras are the reference category (the intercept).

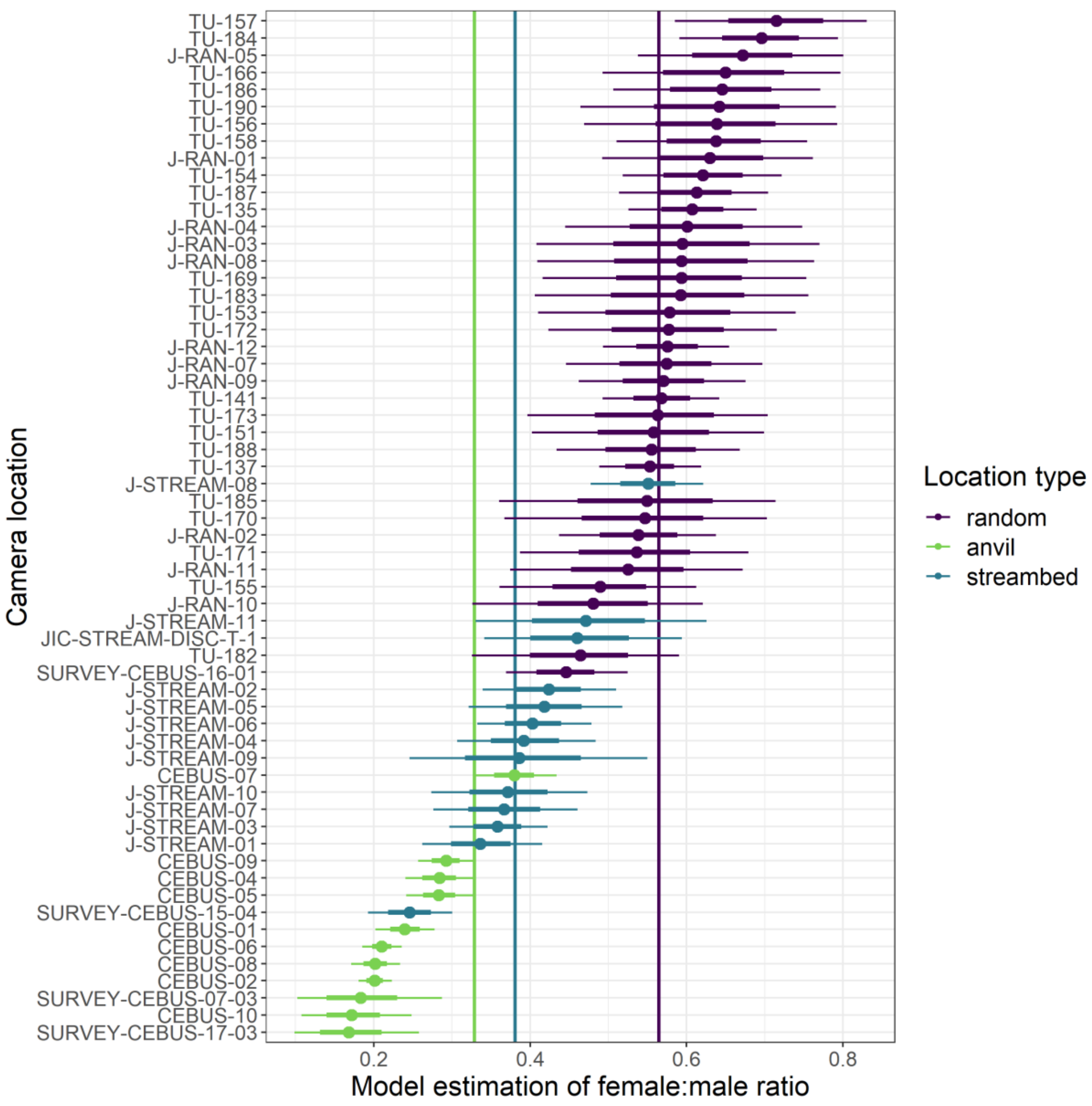


**Figure S1.** Posterior mean estimates of the female:male ratio per camera location on Jicarón. Vertical lines indicate mean estimates per location type (purple for random, green for anvil, blue for streambed). Thick horizontal lines represent 95% credible intervals.

**Model 1b: Estimating adult female:adult male ratio on Coiba**

| | *Estimate* | *CI_95_low* | *CI_95_high* |
|---|---|---|---|
| *Group-Level Effects* | | | |
| Camera location | 0.57 | 0.02 | 1.89 |
| *Population-Level Effects* | | | |
| Intercept | 1.18 | 0.43 | 1.70 |

**Table S4.** Posterior mean model estimates of model estimating sex ratio on streambed cameras on Coiba, a Bayesian logistic mixed model. Model's explanatory power is moderate ($R^2$ = 0.39, 95% CI [0.34, 0.43]). All estimated effects are on logit scale.

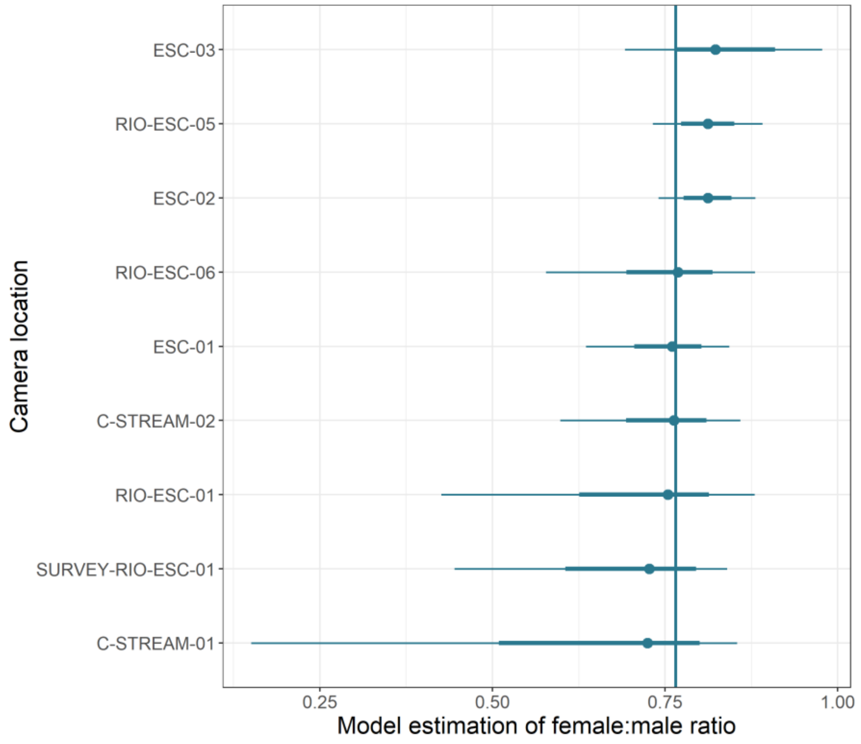


**Figure S2.** Posterior mean estimates of the female:male ratio per camera location. Vertical line indicates the mean estimate. Thick horizontal lines represent 95% credible intervals.

**<u>Model 2: Comparing adult female with infants:adult females without infants ratio between location types on Jicarón</u>**

| | *Estimate* | *CI_95_low* | *CI_95_high* |
|---|---|---|---|
| *Group-Level Effects* | | | |
| Camera location | 0.48 | 0.29 | 0.70 |
| Month | 0.35 | 0.18 | 0.63 |
| *Population-Level Effects* | | | |
| Intercept | -1.34 | -1.76 | -0.93 |
| Location type: Anvil | 0.16 | -0.59 | 0.44 |
| Location type: Streambed | -0.16 | -0.90 | 0.86 |
| Distance to coast | -0.08 | -0.25 | 0.09 |
| Interaction Anvil x Distcoast | -0.52 | -1.89 | 0.97 |
| Interaction Streambed x Distcoast | 0.75 | 0.12 | 1.36 |

**Table S5.** Posterior mean model estimates of model comparing ratio of females with infants:females without infants across location types on Jicarón, a Bayesian logistic mixed model. Model's explanatory power is weak ($R^2$ = 0.13, 95% CI [0.10, 0.16]). All estimated effects are on logit scale. Random cameras are the reference category (the intercept).

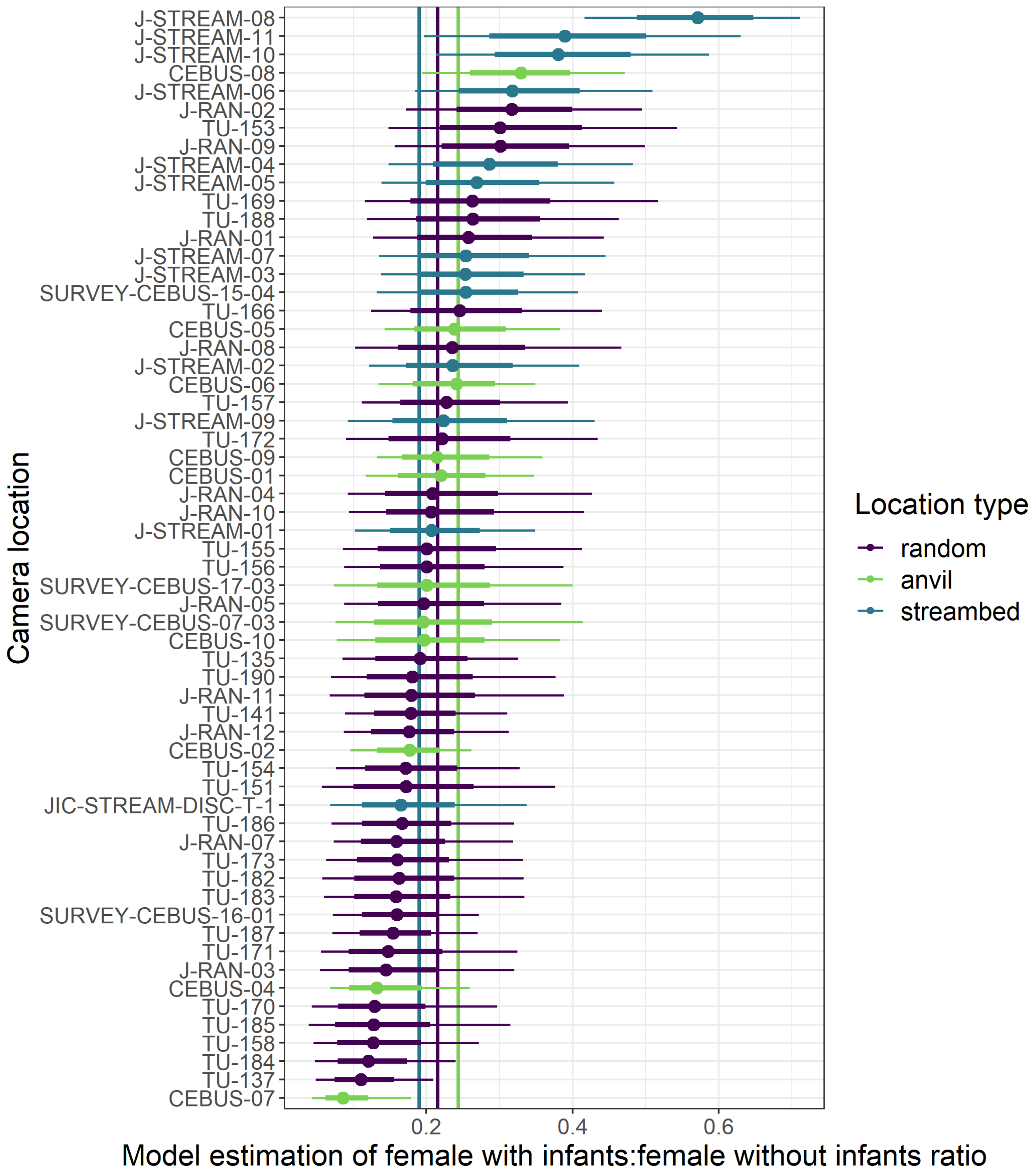


**Figure S3.** Posterior mean estimates of female with infants:female without infants ratio per camera location. Vertical lines indicate mean estimates per location type (purple for random, green for anvil, blue for streambed). Thick horizontal lines represent 95% credible intervals.

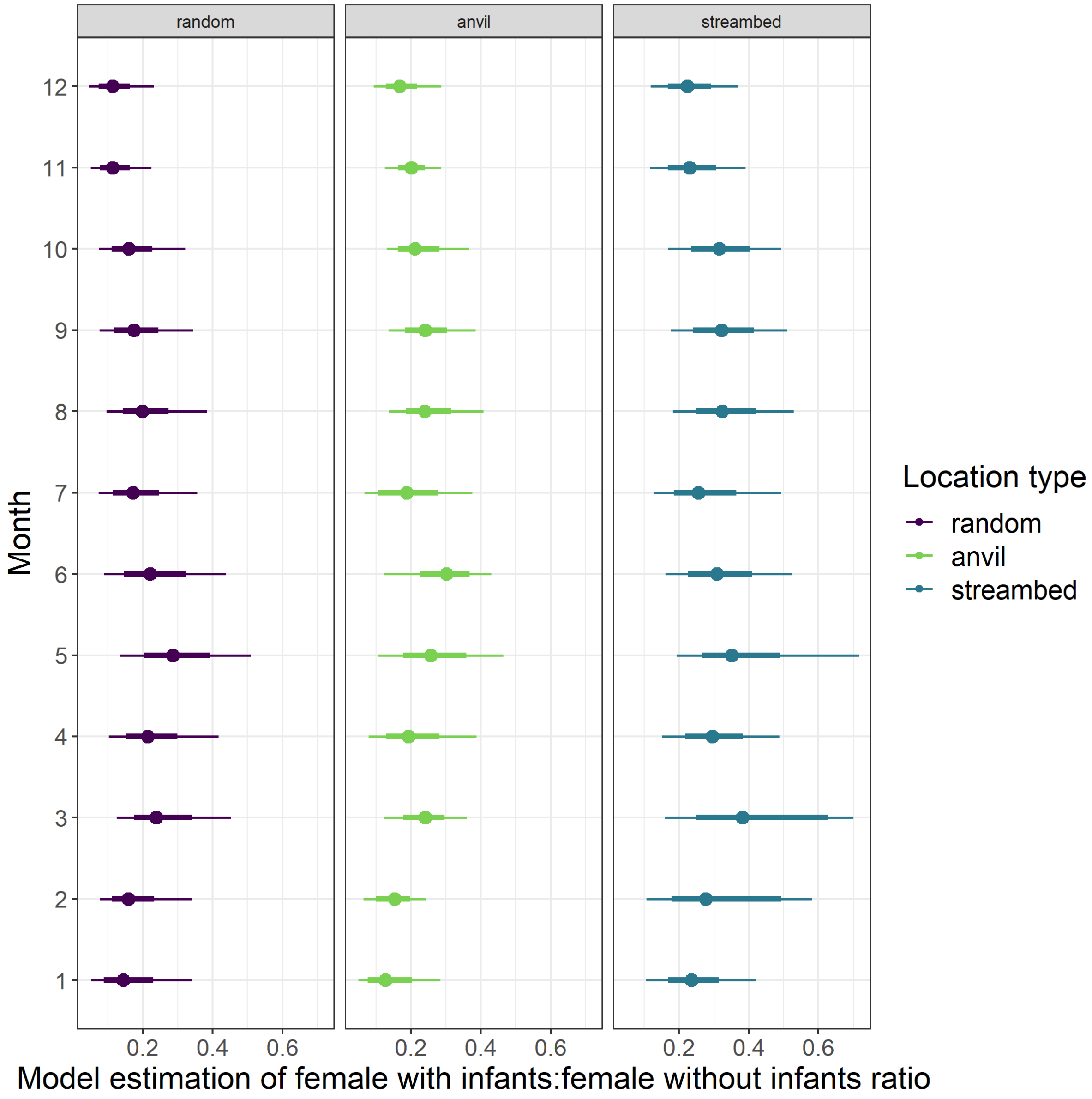


**Figure S4.** Posterior mean estimates of female with infants:female without infants ratio per month and per location type. Thick horizontal lines represent 95% credible intervals.

**Model 2b: Comparing adult female with infants:adult females without infants ratio at streambed cameras on Coiba**

| | *Estimate* | *CI_95_low* | *CI_95_high* |
|---|---|---|---|
| *Group-Level Effects* | | | |
| Camera location | 2.60 | 1.23 | 4.94 |
| *Population-Level Effects* | | | |
| Intercept | -2.47 | -4.67 | -0.55 |

**Table S5.** Posterior mean model estimates of model comparing ratio of females with infants:females without infants across location types on Coiba, a Bayesian logistic mixed model. Model's explanatory power is moderate ($R^2$ = 0.38, 95% CI [0.25, 0.48]). All estimated effects are on logit scale.

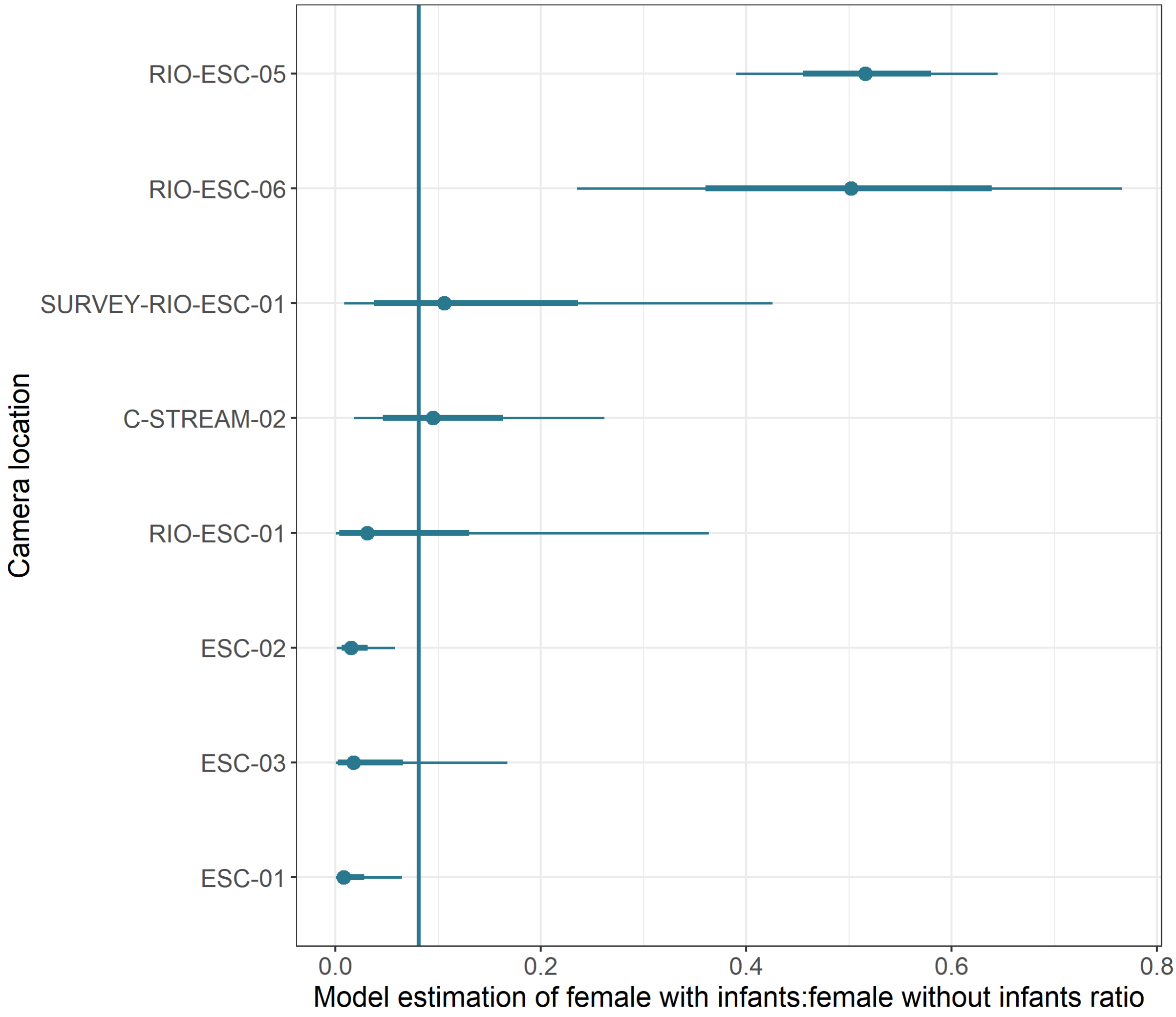


**Figure S5.** Posterior mean estimates of female with infants:female without infants ratio per camera location on Coiba. Vertical line indicates the mean estimate. Thicker horizontal lines represent 95% credible intervals.
